# Cooperativity of membrane-protein and protein-protein interactions control membrane remodeling by epsin 1 and regulate clathrin-mediated endocytosis

**DOI:** 10.1101/2020.01.27.920827

**Authors:** Benjamin Kroppen, Nelli Teske, King F. Yambire, Niels Denkert, Indrani Murkhejee, Daryna Tarasenko, Garima Jaipuria, Markus Zweckstetter, Ira Milosevic, Claudia Steinem, Michael Meinecke

**Affiliations:** Department of Cellular Biochemistry, University Medical Center Göttingen, Humboldtallee 23, 37073 Göttingen, Germany; Institute for Organic and Biomolecular Chemistry, University of Göttingen, Tammannstr. 2, 37077 Göttingen, Germany; European Neuroscience Institute Göttingen – A Joint Initiative of the University Medical Center Göttingen and the Max-Planck-Society, Grisebachstr. 5, 37077 Göttingen, Germany; German Center for Neurodegenerative Diseases (DZNE), Von-Siebold-Str. 3a, 37075 Göttingen, Germany; Max Planck Institute for Biophysical Chemistry, Am Fassberg 11, 37077 Göttingen, Germany; Göttinger Zentrum für Molekulare Biowissenschaften – GZMB, 37077 Göttingen, Germany

**Author notes:** Correspondence to &.

## Abstract

Membrane remodeling is a critical process for many membrane trafficking events, including clathrin-mediated endocytosis. Several molecular mechanisms for protein induced membrane curvature have been described in some detail. Contrary, the effect that the physico-chemical properties of the membrane has on these processes is far less well understood. Here, we show that the membrane binding and curvature-inducing ENTH domain of epsin1 is regulated by phosphatidylserine (PS). ENTH binds to membranes in a PI(4,5)P_2_-dependent manner but only induces curvature in the presence of PS. On PS-containing membranes, the ENTH domain forms rigid homo-oligomers and assembles into clusters. Membrane binding and membrane remodeling can be separated by structure-to-function mutants. Such oligomerization mutants bind to membranes but do not show membrane remodeling activity. In vivo they are not able to rescue defects in epidermal growth factor receptor (EGFR) endocytosis in epsin knock-down cells. Together, these data show that the membrane lipid composition is important for the regulation of protein-dependent membrane deformation during clathrin-mediated endocytosis.

## Introduction

Clathrin-mediated endocytosis (CME) is a special form of vesicle budding from the plasma membrane that capitalizes on a formation of clathrin lattices around maturing buds. In this orchestrated and highly dynamic process some 50 proteins come together to internalize receptors, extracellular ligands or to recycle synaptic vesicles ^1,2^. Over the course of several seconds the plasma membrane undergoes heavy remodeling. A clathrin-coated pit is formed and eventually pinched of the plasma membrane to release a highly curved clathrin-coated vesicle (CCV).

Over the years a number of proteins with the ability to deform membranes were identified to be important in this process ^3,4^. One family of these proteins are epsins. Epsins are evolutionary conserved proteins, which in mammals consist of three genes, epsin 1 – 3 ^5^. Epsin1 consists of a long, unfolded, C-terminal protein-protein interaction domain and an N-terminal membrane binding domain. With the N-terminal ENTH domain (epsin N-terminal homology), epsin1 binds to the cytoplasmic surface of the phosphatidylinositol-4,5-bisphosphate (PI(4,5)P_2_)-containing plasma membrane. Epsin was shown to be involved in CME especially in the uptake of epidermal growth factor (EGF) ^6^. It has also been implicated in CME of other cargos like transferrin but here its role is less well understood ^6,7^. Upon PI(4,5)P_2_ binding a formerly unstructured region at the very N-terminus folds into an amphipathic helix, called helix 0 ^8,9^. This amphipathic helix inserts into the membrane, which probably induces membrane curvature, which is believed to be important to drive clathrin-coated pit formation ^10^. Despite the necessity of PI(4,5)P_2_ to bind the ENTH domain, almost nothing is known about the involvement of the membrane lipid composition in protein-dependent membrane deformation.

In this study we have identified the inner leaflet plasma membrane lipid phosphatidylserine (PS) to be crucial for the membrane curvature inducing activity of the ENTH domain. Presence of this lipid triggers homo-oligomerization of the epsin protein, and through structure-function analysis we show that this assembly is important for membrane deformation.

## Results

### Epsin1 ENTH domain induces phosphatidylserine-dependent membrane curvature

The role of the membrane lipid composition in the process of protein-induced membrane deformation is poorly understood. Due to the complexity and heterogeneity of biological membranes *in vitro* studies usually employ ill-defined lipid mixtures derived from native tissue or very simple compositions that show little resemblance to the *in vivo* situation. Often these two different approaches lead to results that are difficult to compare, or are even contradicting. When analyzing the well-studied curvature-inducing activity of epsin1 ENTH domain, we observed that the ENTH domain is able to bind to large unilamellar vesicles (LUVs) of different membrane composition in a PI(4,5)P_2_-dependent manner, as described before (**Figure 1A**) ^8,11^. Surprisingly though, we found that while addition of the ENTH domain to vesicles made from brain-derived lipid mixtures led to deformed membranes (**Figure 1B-C**), no signs of membrane remodeling was observed in liposomes formed from phosphatidylcholine (PC), phosphatidylethanolamine (PE) and PI(4,5)P_2_ (**Figure 1F**(-PS)). These effects could be visualized by electron microscopy (**Figure 1B & 1F**) and in bulk by analyzing the size distribution of LUVs by dynamic light scattering (**Figure 1C & 1G**).

**Figure 1:**
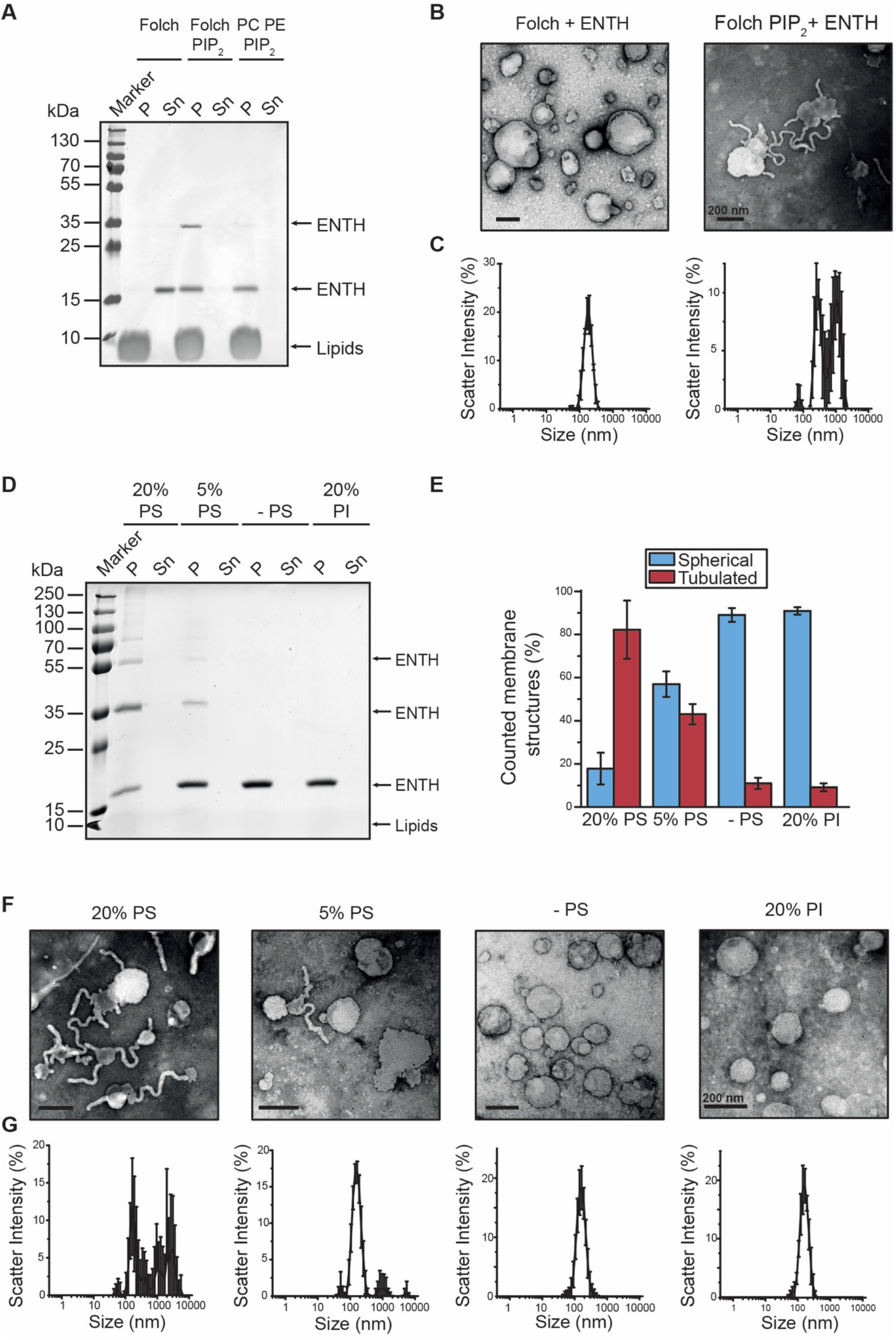
ENTH deforms membranes in a PS-dependent manner. **(A)** ENTH domain binding to LUVs only depends on the presence PI(4,5)P_2_. Protein binding was detected as ENTH domain co-sedimentation along with LUVs in spin assays. The ENTH domain did not bind to LUVs consisting only of a polar brain lipid extract (Folch) but binding was observed if LUVs were supplemented with PI(4,5)P_2_ (95 : 5 mol%). The same effect was observed if LUVs were composed of PC, PE, PI(4,5)P_2_ (65 : 30 : 5 mol%). **(B)** ENTH domain induced membrane deformation is associated to its binding to membranes. As LUVs were exclusively composed of Folch lipids no ENTH domain binding was expected and no membrane deformation was observed in electron microscopy. In contrast, LUVs displayed membrane deformation if LUVs were composed of Folch and PI(4,5)P_2_ (95 : 5 mol%). **(C)** The result was confirmed by DLS showing a single sharp peak if LUVs were composed exclusively of Folch lipids and a diffuse scattering profile if LUVs were composed of Folch and PI(4,5)P_2_ indicating for membrane deformation. DLS data were acquired by 3 times 3 independent measurements, each one consisting of 21 repetitions for each lipid composition. Error bars represent the standard error of the mean (SEM). **(D)** The ENTH domain forms homo-oligomers on LUVs induced by the presence of PS. By increasing the PS concentration within lipid concentrations for LUVs the ENTH-domain oligomer bands became more pronounced at approx. 35, 60 and above 70 kDa. The effect is PS specific because if 20% PI was used instead of 20%PS no ENTH domain oligomer bands appeared. LUVs containing PS were composed of PC, PE, PS and PI(4,5)P_2_ (20% PS – 45 : 30 : 20 : 5 mol%; 5% PS – 60 : 30 : 5 : 5 mol%). LUVs without PS were composed of PC, PE and PI(4,5)P_2_ (0% PS – 65 : 30 : 5 mol%) and LUVs containing PI were composed of PC, PE, PI, PI(4,5)P_2_ (20% PI – 45 : 30 : 20 : 5 mol%). **(E)** The statistical evaluation of EM micrographs displays that if LUVs containing 20% PS approx. 85% of counted membrane structure were deformed. With decreasing PS concentration within LUVs deformed membranous structures became less predominant and with 5% PS approx. 45% of counted membranous structures were tubulated. Without PS and if 20% PI was used instead of PS only a small amount of membranous structures were deformed (approx. 10%) **(F)** PS promotes ENTH domain induced membrane deformation of LUVs with a defined lipid composition. The electron microscopy of LUVs showed that with increasing concentrations of PS the ENTH domain induced membrane deformation becomes more pronounced and no membrane deformation was observed in absence of PS or if 20% PI was added instead. LUVs were composed with similar lipid compositions as in Figure 1D. At least 250 membrane structures were counted for each condition and the error bars represent the standard deviation. **(G)** DLS analysis confirmed the results with LUVs containing 20% PS caused a highly diffuse scattering profile indicating for massive membrane deformation. Decreasing the PS concentration resulted in a less diffuse scattering profile and in absence of PS (- PS) and if LUVs were supplemented with PI (20% PI) a sharp peak was observed. LUVs were composed with similar lipid compositions as in Figure 1D and the DLS data was acquired by 3 independent measurements, each one consisting of 21 repetitions for each lipid composition. Each measuring circle was repeated 3 times.

PS is the most abundant anionic cellular phospholipid. In resting cells, it is with at least 15 % exclusively found in the inner leaflet of the plasma membrane ^12^. We hence asked if the presence of PS would influence membrane deformation by the ENTH domain. The ENTH domain of epsin1 efficiently bound to LUVs consisting of PC, PE, PI(4,5)P_2_ and increasing amounts of PS (**Figure 1D**). Membrane deformation was dependent on the presence of PS and statistical evaluation of electron micrographs of LUVs revealed a clear correlation between PS concentration and membrane remodeling (**Figure 1E-F**). Similar results were obtained using giant unilamellar vesicles (GUVs) (**Figure S1A-B**). Membrane curvature induction was specific to PS as the presence of another anionic lipid, phosphatidylinositol (PI), did not lead to deformed membranes (**Figure 1E-G**). High concentrations of unsaturated PE also did not lead to membrane deformation, ruling out that changes of membrane morphology are induced by high concentration of non-bilayer lipids (**Figure S1C**).

### PS induces ENTH domain oligomerization

The liposome co-sedimentation assays showed that the ENTH domain of epsin1 binds to membranes in a PI(4,5)P_2_-dependent manner, as published before (**Figure 1A**). The amount of bound protein did not vary with changing lipid compositions (**Figure 1A & 1D**). However, when analyzing the fractions of liposome bound proteins, SDS resistant oligomers were detected when using Folch liposomes (**Figure 1A**). The same tendency for oligomerization was observed with increasing PS concentration (**Figure 1D & 2A**). As the appearance of PS-dependent oligomers coincided with membrane deformation, we analyzed this effect in more detail. We first tested if oligomerization not only occurs in SDS resolved samples but can also take place on membranes. We therefore performed FRET (Förster resonance energy transfer) experiments of ENTH domain in the presence of PS-free and PS-containing LUVs. Efficient FRET signals were detected when two different fractions of fluorescently labeled protein (ENTH-Atto-488 [donor] and ENTH-Atto-532 [acceptor]) were incubated with PS-containing liposomes (**Figure 2B**). PS-free LUVs led to significantly reduced FRET signals, confirming that ENTH molecules are in close proximity on PS-containing membranes. Interestingly, when repeating membrane binding and membrane deformation with PI(4,5)P_2_-containing liposomes and ENTH domains, we observed that the presence of the PS head-group was sufficient to partially restore oligomerization and membrane remodeling (**Figure 2C-F**).

**Figure 2:**
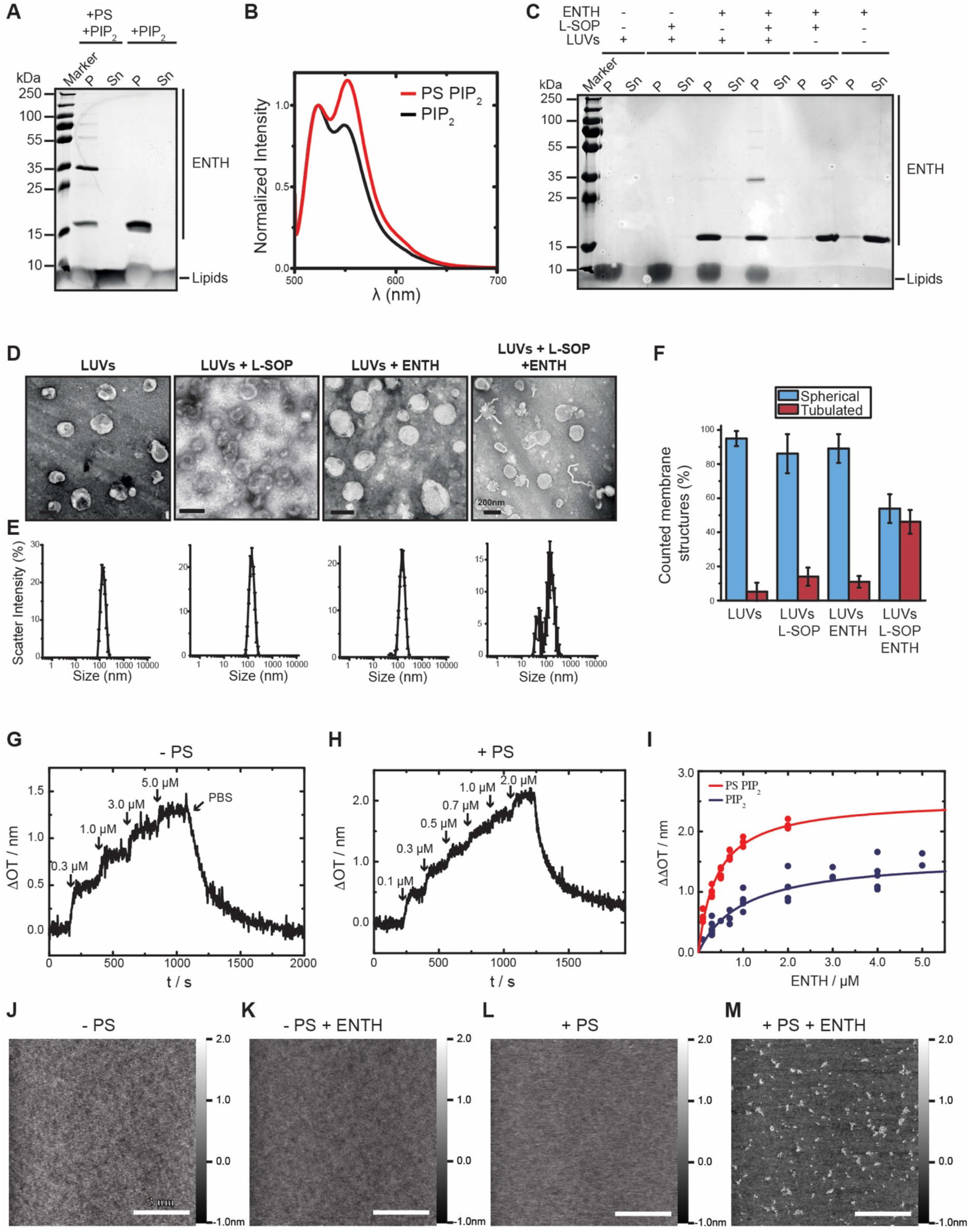
PS-dependent oligomerization of ENTH. **(A)** In spin assays the co-sedimentation of the ENTH domain was accompanied by the formation of homo-oligomers in presence of PS. If LUVs did not contain PS the ENTH domain still co-sedimented but no oligomerization was observed. LUVs were composed of PC, PE, PS and PI(4,5)P_2_ (PS PIP_2_ - 45 : 30 : 20 : 5 mol%) or PC, PE and PI(4,5)P_2_ (PIP_2_ - 65 : 30 : 5 mol%). **(B)** Protein-protein interaction was demonstrated by FRET assay in which the normalized spectra showed that the ENTH domain displayed a FRET effect if LUVs contained PS (increase of acceptor emission at 552 nm). In comparison, decreased acceptor emission was observed if we used LUVs without PS. For FRET experiments two differently labeled ENTH domain populations were generated, the one was labeled with Atto488 maleimid, the other population was labeled with Atto532 maleimid. LUVs were composed as described in (A). **(C)** The soluble headgroup of PS O-phosphoserine (L-SOP) is able to promote the homo-oligomerization of the ENTH domain on LUVs lacking PS. LUVs incubated only with L-SOP behaved the same way as if L-SOP was omitted. The ENTH domain co-sedimented along with LUVs in absence of L-SOP, but no oligomerization pattern was observed. In contrast together with L-SOP the ENTH domain formed homo-oligomers on LUVs. If the ENTH domain was not incubated with LUVs no homo-oligomers were observed in presence and absence of L-SOP. L-SOP was used in a 4 times molar excess to the ENTH domain. LUVs were composed of PC, PE and PI(4,5)P_2_ (PIP_2_ - 65 : 30 : 5 mol%). **(D)** L-SOP also recued partially the ENTH domain dependent membrane deformation on LUVs. L-SOP alone and the ENTH domain alone did not have any effects on LUVs that remained undeformed. If LUVs were incubated together with L-SOP and the ENTH domain a membrane deformation effect was observed, but not to the extent if as PS was added to the LUV lipid composition. L-SOP was used in a 4 times molar excess to the ENTH domain and LUVs were composed of PC, PE and PI(4,5)P_2_ (PIP_2_ - 65 : 30 : 5 mol%). Scale bars correspond to 200 nm. **(E)** ENTH-domain dependent membrane deformation in presence of L-SOP on LUVs without PS was confirmed by DLS experiments. DLS peaks of LUVs alone, LUVs incubated with L-SOP or ENTH domain alone were monodisperse and did not indicate membrane deformation. If LUVs were incubated together with ENTH-domain and L-SOP the scattering profile was heterogenic indicating membrane deformation. L-SOP and LUVs were prepared as described before (D). DLS data were acquired by 3 times 3 independent measurements, each one consisting of 21 repetitions for each lipid composition. Error bars represent the standard error of the mean (SEM). **(F)** The statistical evaluation of the micrographs displays that LUVs incubated L-SOP or ENTH domain alone did not induce membrane deformation. But if LUVs were incubated with L-SOP and ENTH domain together about 50% of the counted membrane structure were tubulated. At least 150 membrane structures were counted. The error bars were calculated by the standard deviation. **(G)** Adsorption of ENTH^WT^ to a POPC/PI(4,5)P_2_ (95:5) bilayer. The stepwise increase in Δ*OT* upon addition of different ENTH concentrations (marked by arrows) shows the specific binding of ENTH to PI(4,5)P_2_. Rinsing with PBS results in desorption of the protein indicating reversible binding. **(H)** Time-resolved change in optical thickness upon addition of different ENTH^WT^ concentrations (marked by arrows) to a POPC/POPS/PI(4,5)P_2_ (75:20:5) bilayer. After rinsing with PBS, almost all bound protein desorbs from the membrane indicating reversible binding. **(I)** Adsorption isotherms of ENTH^WT^ to POPC/PI(4,5)P_2_ (95:5) (blue circles) and POPC/POPS/PI(4,5)P_2_ (75:20:5) (red circles) bilayers. A Langmuir adsorption isotherm was fit to the data (solid lines). Non-linear regression weighted by the corresponding number of measurements that went into each concentration was carried out using a Levenberg-Marquardt algorithm. The obtained values for ΔΔ*OT*_max_ and *K*_D_ are summarized in Tab. 1. **(J-M)** Atomic force micrographs of (J) DOPC/DOPE/PI(4,5)P_2_/Texas Red DHPE (64.9/30/5/0.1) and (L) DOPC/DOPE/DOPS/PI(4,5)P_2_/Texas Red DHPE (44.9/30/20/0.1) bilayers on hydrophilic silicon dioxide wafers prior ENTH addition. (K) and (M) show the corresponding micrographs after 2 h of ENTH^WT^ incubation (1 μM). Only in the presence of DOPS, protein clusters were observed on the membrane surface. From the topography images, the occupancy of 6 ± 1 % and the protein height of 1.2 ± 0.2 nm (values ± SD, *n* = 32, with *n* the number of evaluated images from three independent experiments) were calculated.

**Table 1:** Summary of the fit results of the Langmuir adsorption isotherms for ENTH^WT^ and ENTH^R114A^. The values are given as parameter ± SE.

|  | ***K<sub>d</sub>*** |  | **$\Delta\Delta OT_{max}$** |  |
| --- | --- | --- | --- | --- |
|  | ENTH<sup>WT</sup> | ENTH<sup>R114A</sup> | ENTH<sup>WT</sup> | ENTH<sup>R114A</sup> |
| **no PS** | 1.0 ± 0.2 μM | 2.9 ± 0.8 μM | 1.6 ± 0.1 nm | 1.5 ± 0.2 nm |
| **with PS** | 0.42 ± 0.05 μM | 1.0 ± 0.3 μM | 2.5 ± 0.1 nm | 2.7 ± 0.2 nm |

To understand the molecular details of this lipid-specific protein-membrane interaction, we next quantitatively investigated the behavior of the ENTH domain on different membranes. No signs of ENTH domain membrane binding were observed in the absence of PI(4,5)P_2_ by reflectometric interference spectroscopy (RIfS) (**Figure S2B**). When the ENTH domain was added to supported bilayers containing PI(4,5)P_2_ an increase in height of about 0.6 nm was observed (**Figure 2G**). When PS was included in these experiments, the increase was significantly higher (**Figure 2H**). Plotting the change in optical thickness against the ENTH domain concentrations allowed us to calculate binding affinities for membranes of different composition. The dissociation constant *K*_D_ for PI(4,5)P_2_ containing PC membranes was about 3 μM. *K*_D_ decreased to about 0.4 μM when the membranes contained 20 % PS (**Figure 2I**), indicating a higher binding affinity of ENTH to PS-containing membranes. We then sought to analyze structural details of the PS-dependent oligomer formation and higher protein affinity. When the ENTH domain was added to PI(4,5)P_2_-containing bilayers in the absence / presence of PS and the membrane surface was analyzed by atomic force microscopy (AFM) clear protein clusters could be observed in a PS-dependent manner (**Figure 2K-M**). Whereas membranes of various composition in the absence of proteins showed a rather smooth surface, the addition of ENTH domain led to protein clusters or oligomers in line with the biochemical and FRET results. Control experiments with PS-free membranes, in which no signs of protein clusters were observed (**Figure 2K**), support the idea of PS-dependent ENTH oligomerization.

To further analyze the structural differences of membrane bound ENTH domains in the absence of presence of PS, we used solid-state NMR spectroscopy. The ENTH domain was ^13^C/^15^N-labeled by expressing the protein in *E. coli* cells grown in buffer supplemented with ^15^N ammonium chloride and ^13^C glucose. Though the measured solid-state NMR spectra were too inhomogeneous to reach atomic resolution, comparison of 1D ^13^C-spectra showed clear differences between the presence of PS, where ENTH is able to tubulate membranes, and the absence of PS (**Figure S2C-F**). ^1^H-^13^C CP experiments revealed an increased signal intensity in C^α^ and C^β^ regions in the presence of PS (**Figure S2E**), which is an indicator of increased rigidity of the protein. A rigid protein population with a reduced number of motions supports the results showing PS-dependent oligomerization in biochemical, FRET and AFM experiments. It also explains results from AFM measurements in the absence of PS. The relatively even surface observed in these experiments is indicative of mobile protein monomers that cannot be resolved by AFM (**Figure 2K**).

### A single amino acid exchange abolishes ENTH domain oligomerization and membrane deformation

It is well-studied that upon binding to PI(4,5)P_2_ an unstructured N-terminal stretch of the ENTH domain folds into an amphipathic α-helix that inserts into the membrane ^13,14^. It is however not obvious how this mode of binding allows the observed PS-dependency. In models of the membrane bound ENTH domain based on EPR measurements a self-assembly was suggested, and beside the hydrophobic PI(4,5)P_2_ binding pocket and the amphipathic helix 0 a loop region was found in close proximity to the membrane ^13,15^. Interestingly, an arginine residue in position 114 is located at the tip of this loop (**Figure 3A**).

**Figure 3:**
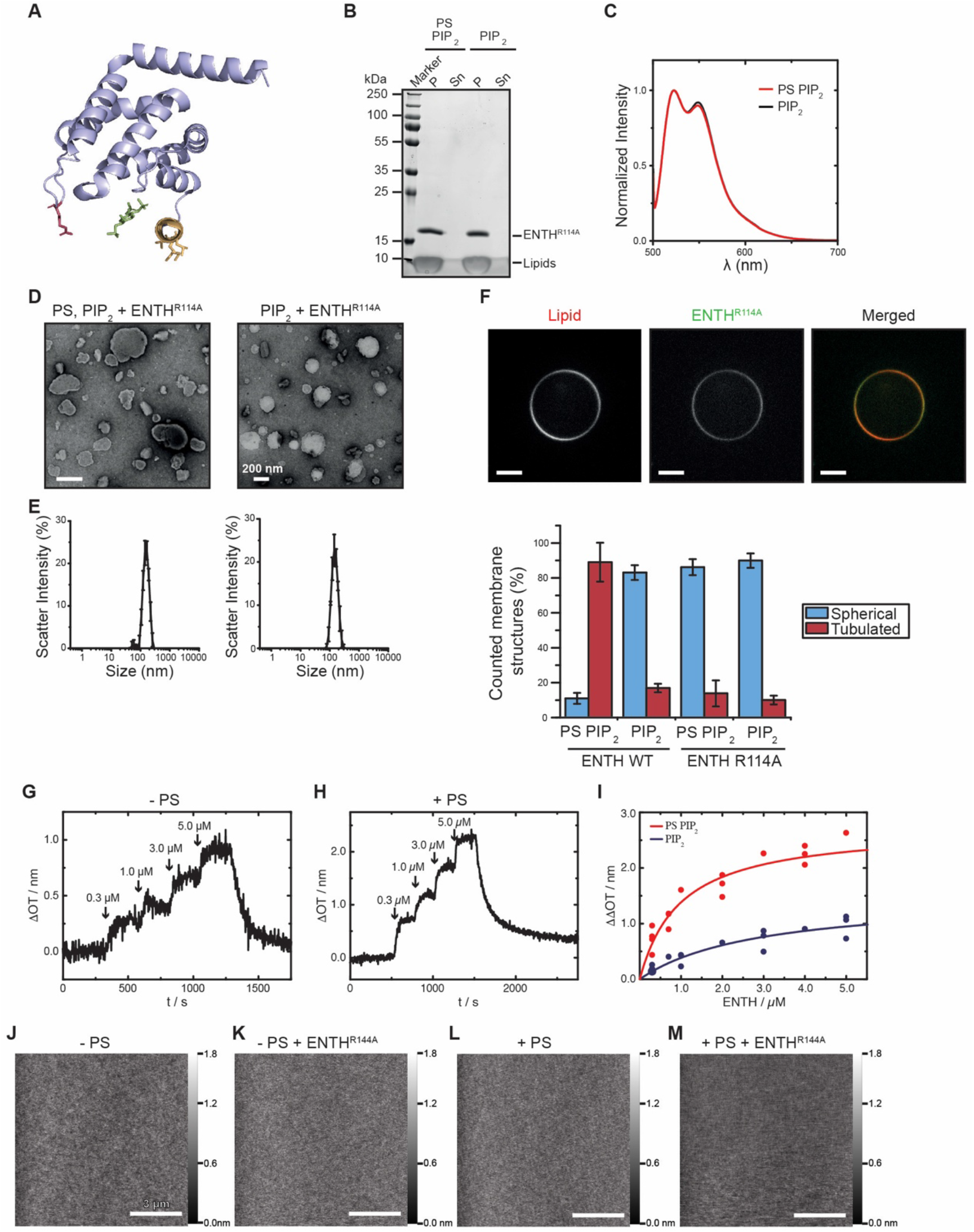
A point mutation in ENTH eliminates PS-dependent oligomerization and curvature induction. **(A)** Arginine 114 (R114) is located within the loop l6,7 in the crystal structure of the ENTH domain. The amphipathic helix (yellow) structures upon interaction with the PI(4,5)P_2_ headgroup inositol (1,4,5)-triphosphate (green). This arginine was mutated to alanine by site-directed mutagenesis (ENTH^R114A^). **(B)** Similar to the wild type protein ENTH domain R114A co-sediments with LUVs containing PI(4,5)P_2_ in spin assays. But in contrast to the wild type ENTH domain no formation of oligomers were observed in presence of PS. LUVs used for this spin assay were composed of PC, PE, PS, PI(4,5)P_2_ (“PS PIP_2_” - PC 45 : 30 : 20 : 5 mol%; “PIP_2_” – 65 : 30 : 5 mol%) **(C)** ENTH^R114A^ does not form homo-oligomers after being recruited to membranes during protein FRET experiments. Two differently labeled protein populations were generated, one labeled with Atto532, the other one labeled with Atto488. LUVs for the experiment were composed of PC, PE, PS and PI(4,5)P_2_ : (“PS PIP_2_” - 45 : 30 : 20 : 5)mol%; PIP_2_ – 65 : 30 : 5 mol%). **(D)** ENTH^R114A^ is not able to induce membrane deformations on LUVs containing PS like the ENTH domain WT protein and no membrane tubules were observed during electron microscopy. LUVs were composed as described before (see 3B) and scale bars correspond to 200 nm. **(E)** The corresponding DLS profiles confirm the results and only a sharp peak appeared in absence and presence of PS within lipid compositions indicating for no membrane deformation. LUVs were composed as described before (see 3B). DLS data were acquired by 3 times 3 independent measurements, each one consisting of 21 repetitions for each lipid composition. Error bars represent the standard error of the mean (SEM). **(F)** ENTH^R114A^ binds to membranes but fails to induce membrane deformation also in GUV experiments. After incubation the protein signal (Atto488, green) completely co-localizes at the GUV membrane (Atto847N, red) but no deformation or defined protein induced GUV collapse was observed. GUVs were composed of PC, PE, PS, PI(4,5)P_2_ and Atto647N DOPE (44.5 : 30 : 20 : 5 : 0.5 mol%). ENTH^R114A^ was labeled with Atto488 maleimide by cysteine modification with an DOL of 1.1. The scale bars correspond to 8 μm. **(G)** Statistical analysis of micrographs by ENTH^R114A^ in comparison to the ENTH^WT^. The statistics revealed only the ENTH domain WT deformed membranes if the LUVs contained PS and PI(4,5)P2 and approx. 90% of the counted membranous structures were tubulated. In comparison, LUVs without PS remained undeformed. ENTH^R114A^ did not show membrane deformation effects on LUVs in all tested conditions. At least 200 membrane structures were counted for each used condition, error bars represent the standard deviation. **(H)** Adsorption of ENTH^R114A^ to a POPC/PI(4,5)P_2_ (95:5) bilayer. Upon addition of different ENTH^R114A^ concentrations (marked by arrows) a stepwise increase in Δ*OT* occurs showing the specific binding of ENTH^R114A^ to PI(4,5)P_2_. After rinsing with PBS, the protein desorbs from the membrane indicating reversible binding. **(I)** Time-resolved change in optical thickness upon addition of different ENTH^R114A^ concentrations (marked by arrows) to a POPC/POPS/PI(4,5)P_2_ (75:20:5) bilayer. The protein desorbs after rinsing with PBS showing the reversibility of binding. **(J)** Adsorption isotherms of ENTH^R114A^ to POPC/PI(4,5)P_2_ (95:5) (blue circles) and POPC/POPS/PI(4,5)P_2_ (75:20:5) (red circles) bilayers. The values for ΔΔ*OT*_max_ and *K*_D_ (Tab. 1) were obtained by fitting a Langmuir adsorption isotherm (solid lines) to the data. Non-linear regression weighted by the corresponding number of measurements that went into each concentration was carried out using a Levenberg-Marquardt algorithm. **(K-N)** Atomic force micrographs of (K) DOPC/DOPE/PI(4,5)P_2_/Texas Red DHPE (64.9/30/5/0.1) and (M) DOPC/DOPE/DOPS/PI(4,5)P_2_/Texas Red DHPE (44.9/30/20/0.1) bilayers on hydrophilic silicon dioxide wafers prior ENTH addition. The corresponding micrographs (L, N) were obtained after 2 h of ENTH^R114A^ incubation (1 μM). Even in the presence of DOPS, no protein clusters were observed on the membrane surface indicating the necessity of the amino acid R114 for ENTH cluster formation.

Having found a regulatory effect of the anionic phospholipid PS we next asked the question if membrane binding and / or membrane remodeling of the ENTH could be influenced by this amino acid residue. ENTH domains with a single amino acid exchange from arginine to alanine (R114A) were constructed and analyzed in a similar fashion as the wild-type (WT) protein. The mutation did not affect the overall structure of the ENTH domain as secondary structures of WT and mutant proteins showed no differences when measured by circular dichroism (CD) spectroscopy (**Figure S3A & S3B**). Membrane binding was again observed in a strictly PI(4,5)P_2_-dependent manner. Importantly though, even increasing amounts of PS in LUVs did not lead to SDS resistant oligomers that were observed with ENTH^WT^ (**Figure 3B**). Analysis of differentially labelled membrane bound ENTH^R114A^ showed no PS-dependent differences in FRET efficiency underlining that the mutant fails to form homo-oligomers (**Figure 3C**). When the same vesicles in absence and presence of PS were incubated with ENTH^R114A^ and analyzed for membrane deformation by electron microscopy and dynamic light scattering, no signs of membrane remodeling were observed despite high concentrations of PS (**Figure 3D-F**). Similarly, no membrane curvature induction was observed when GUVs were incubated with ENTH^R114A^ (**Figure S3C**). RIfS measurements with ENTH^R114A^ showed that the mutant is able to bind with similar affinities as the wild-type protein to membranes in the absence and presence of PS (**Figure 3G-H**). Nonetheless, when analyzed by AFM (**Figure 3J-M**) the mutant protein exhibited no sign of PS-dependent cluster formation as the wild-type protein did (**Figure 3M**). When ENTHR114A was analyzed by solid-state NMR spectroscopy no differences in the spectra were detected between the protein bound to a PS-free or PS-containing membrane (**Figure S3C-F**). Taken together, these results clearly demonstrate that PS induced oligomerization lead to membrane deformation, which is inhibited by a point mutation of the ENTH domain.

### Oligomer-dependent membrane deformation is important for clathrin-mediated endocytosis

As we identified a mutant with a clear phenotype in *in vitro* investigations, we next asked if oligomer-dependent membrane remodeling is of physiological importance in clathrin-mediated endocytosis. To this end we performed fluorescent transferrin (Tf) and Epidermal Growth Factor (EGF) uptake assays in Hela cells that were shown before to depend on epsin1 ^6,7^. Specifically, simultaneous depletion of epsin1, 2, and 3 resulted in a significant decrease in Tf uptake (**Figure S4A-C**), in agreement with Boucrot et al. (2012). This perturbation was shown to be specific to CME (Boucrot et al., 2012), which was also confirmed in our experiments where inhibition of CME by clathrin inhibitor Pitstop-2 resulted in a similar decrease in the Tf uptake as the knock-down (KD) of epsin 1, 2 and 3 (**Figure S4B**). Expression of either epsin1-WT-EGFP and epsin1^R114A^-EGFP in epsin 1,2,3, KD HeLa cells was able to restore Tf-A548 uptake to similar levels as seen by flow cytometry (**Figure4A-B**). Curiously, when EGF-Texas Red uptake was examined a different result was obtained: epsin 1,2,3, KD HeLa expressing epsin1^R114A^-EGFP were only able to internalized less fluorescent EGF than cells expressing epsin1^R114A^ -EGFP under the same conditions (**Figure 4C-D**). A similar result was observed when EGF-Texas Red uptake by epsin 1,2,3 KD HeLa expressing epsin1-WT-EGFP and epsin1^R114A^ -EGFP, respectively, was analyzed by fluorescence imaging (**Figure 4E-F**). In sum, our uptake experiments revealed that epsin1^R114A^ alters the EGF uptake, but not the Tf uptake. This is in good agreement with recent reports that EGF-receptor internalization by CME is strictly dependent on epsin, especially in the absence of the adaptor complex AP-2 ^16^. It also underlines earlier observations that knockdown of epsin1 inhibits EGF-receptor internalization while having little effect on endocytosis of transferrin receptors ^6^.

**Figure 4:**
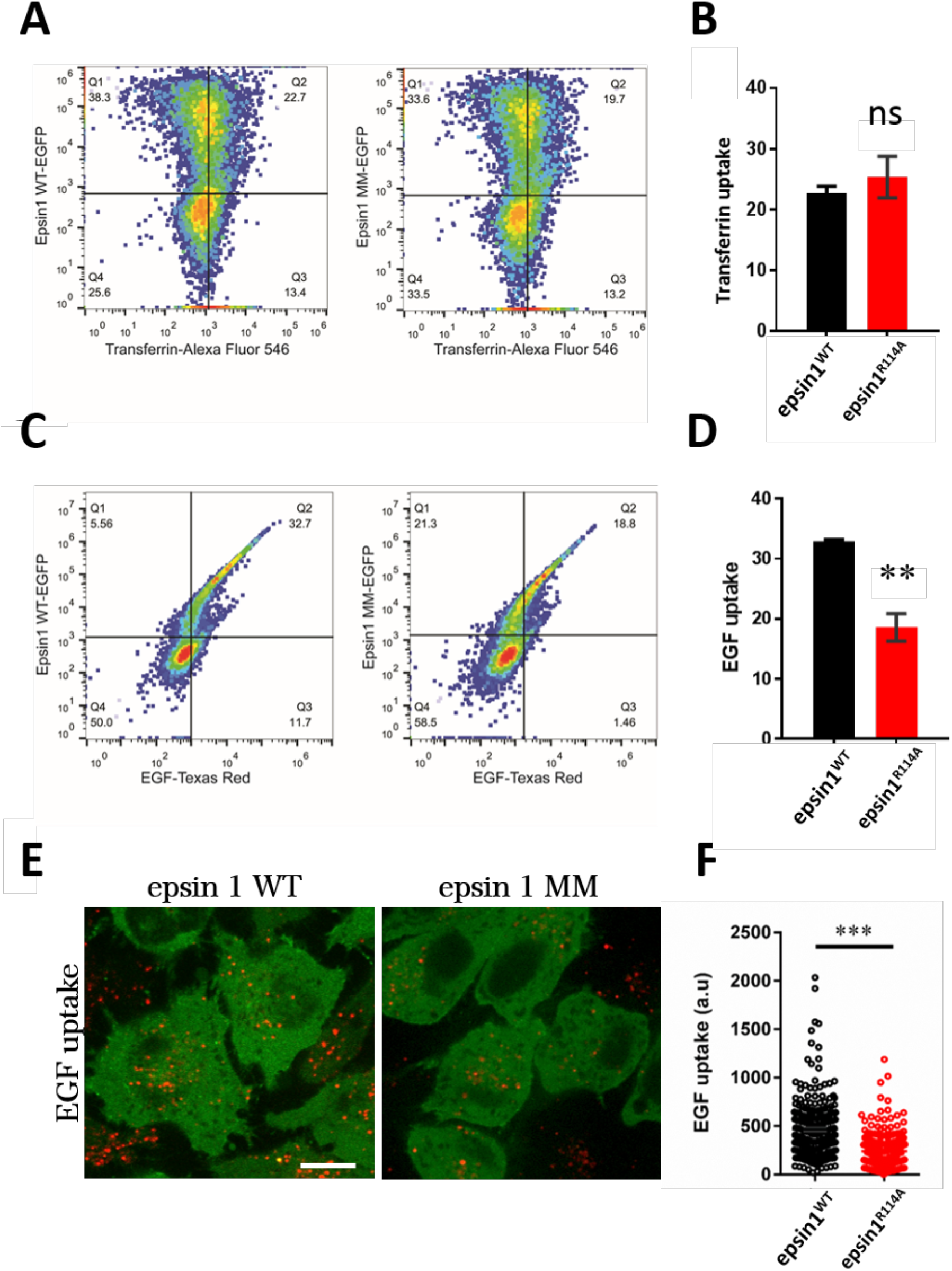
Epsin 1^R114A^ affects the EGF, but not transferrin uptake by HeLa cells. **(A-B)** Epsin 1,2,3 KD HeLa cells expressing either epsin1^WT^-EGFP and epsin1^R114A^-EGFP were allowed to internalize transferrin-Alexa Fluor 548 and were subsequently analyzed by flow cytometry. Experiments were performed as detailed in Methods. 20,000 cells were counted per condition and experiment; four independent experiments were performed. mean ± SEM; ns = not significant (unpaired t-test). **(C-D)** Epsin 1,2,3 KD HeLa cells expressing either epsin1^WT^-EGFP and epsin1^R114A^-EGFP were allowed to internalize EGF-Texas Red and were subsequently analyzed by flow cytometry. Experiments were performed as detailed in Methods. 20,000 cells were counted per condition and experiment; four independent experiments were performed. mean ± SEM; *\*\*p*=0.007 (unpaired t-test). **(E)** Representative images of epsin 1,2,3, KD HeLa cells expressing epsin1^WT^-EGFP and epsin1^R114A^-EGFP after EGF-Texas Red uptake. Although not further analyzed, note that non-transfected (non-fluorescent in green channel) cells in both conditions showed slightly higher levels of internalized EGF-Texas Red. **(F)** Analysis of fluorescent EGF-Texas Red signal in epsin1^WT^-EGFP (black) or epsin1^R114A^-EGFP (red) cells. Data are presented as mean ± SD from three independent experiments. The number of analyzed cells from three consecutive experiments: 226 for epsin1^WT^ and 247 for epsin1^R114A^; ***p<0.001 (unpaired t-test).

## Discussion

By the example of the ENTH domain of epsin1, we herein present the identification of a lipid-dependency for protein-induced membrane remodeling during CME. By combining *in vitro* reconstitution and quantitative biophysical analysis with cell biological approaches we show that PS has a regulatory effect on epsin1. The protein oligomerizes and induces changes in membrane morphology in a PS-dependent manner. PS was implicated to be important for CME ^17,18^. An increased PS concentration in the plasma membrane leads to increased endocytosis rates ^19^ and the proximity of anionic phospholipids may facilitate membrane remodelling ^20^. Our results are, to the best of our knowledge, the first molecular explanation for these results. They also touch on a neglected topic in membrane trafficking - the regulatory involvement of the membrane lipid composition on these processes. As noted before, many proteins operating in CME do not act separately but show cooperative behavior that is most probably necessary for the spatial and temporal fidelity for CME ^21–24^. We now show that the action of proteins during CME is not only fine-tuned by mere membrane recruitment and regulatory protein-protein interactions, but that the membrane lipid composition also plays a crucial role in the regulation of protein-dependent membrane remodelling. This is all the more important as it is still poorly understood why particular proteins act on specific membranes, while being in principle able to bind to other cellular membranes, too. Due to the immense variety and dynamics of cellular lipids, a detailed molecular characterization of the regulatory effect of the membrane composition on membrane-based processes is still missing to a large degree. Nonetheless, sporadic reports are emerging and the importance of such interactions will be in all likelihood widespread.

Clathrin-mediated endocytosis is a process that depends on the spatial and temporal precise action of a complex set of different proteins. These include not only clathrin itself, but also adaptor proteins. Together these proteins mediate cargo sorting, nucleation of the clathrin coated pit, membrane remodeling during pit maturation and fission as well as release of the clathrin-coated vesicle. Besides being heavily studied for its physiological importance for cell signaling, nutrient uptake, migration or neurotransmission, to name just a few, CME has become a model system to study membrane remodeling proteins ^1^. The vast majority of these proteins are peripheral, temporarily attached membrane proteins. They can be expressed and purified as soluble molecules making them ideal candidates for *in vitro*, biochemical and structural investigations. In the case of the ENTH domain of epsins, the molecular mechanism that leads to protein-induced membrane deformation was widely accepted to depend on the membrane insertion of an amphipathic α-helix (helix 0) that folds upon interaction with PI(4,5)P_2_ ^8,9^. Though many reports are in good agreement with this model, a different explanation for the induction of membrane curvature was recently reported that relies on macromolecular crowding ^25,26^. Our own recent results showed that binding of the ENTH domain helix 0 to membranes results in a decrease in lateral membrane tension ^27,28^, which in turn would lead to a decreased membrane bending modulus and results in a membrane more susceptible to deformation. Together with the results presented here it seems reasonable to assume that not only amphipathic helix insertion is necessary for curvature induction but also a tight spatial arrangement of the protein on the membrane’s surface, seen here as oligomerization, contributes to membrane deformation. This is in line with the notion that the ENTH domain together with the related ANTH domain co-assemble into oligomers on membranes ^29,30^.

## Acknowledgements

We thank Catia Diogo and Nuno Raimundo for the help with qPCR, and Rafael Rinaldi Ferreira for the help with FACS. This work was supported by the Deutsche Forschungsgemeinschaft: SFB803 projects A11 (to MZ), B04 (to NT and CS) B09 (to BK and MG), FOR2848 project P5 (to ND and MM) and Emmy Noether Young Investigator Award MI-1702/1 (to IM).

## Methods

### Protein Biochemistry

#### Protein expression and purification

The ENTH domain and its mutants were recombinantly expressed in *E. coli* BL21(DE3) cells (Stratagene, California, USA). Therefore, the cells were transformed by heat shock transformation with pGEX-6P-2 vectors (including genes for Rat Epsin1 WT ENTH 1-164 and generated mutants). The recombinant expression was performed by isopropyl-β-D-thiogalactopyranosid (IPTG) induction. Transformed, selected, single colonies of *E. coli* BL21(DE3) were transferred into LB medium (10 g/l NaCl, 5 g/l yeast extract, 10 g/l tryptone, g/l ampicillin) and cultivated for about 2.5 h at 37°C and 140 rpm agitation until reaching an optical density (OD_600_) of 0.6 - 0.8. The expression culture was substituted with 1 mM IPTG and incubated for 3 h at 30°C and 140 rpm agitation. The expression culture was finally centrifuged for 10 min at 4°C and 5316 × *g* (Thermo Scientific H-12000 BioProcessing Rotor). The cell pellet was separated from the supernatant and resuspended with 40 ml HEPES-buffer per 2 l culture on ice. The cell suspension was subsequently centrifuged for 30 min at 4°C and 3250 × *g* (Eppendorf Swing-bucket rotor A-4-62). The supernatant was discarded. To obtain single-band pure ENTH-domain a three-step chromatographic purification strategy was performed. In the first step the ENTH-domain was isolated from the soluble cell lysate as GST fusion protein by using GSTrapFF™ 5 ml columns together with the ÄKTAPrime Plus system (GE Healthcare Life Science, Chalfont St Giles, UK). The cell lysate was injected onto the column (0.4 ml/min, 8°C) and non-binding fractions were separated by washing with 25 ml GST binding buffer (5 ml/min, 8°C). The target protein was eluted with GST elution buffer (1 ml/min, 8°C). The GST-tag was proteolytically cleaved from the ENTH-domain by using PreScission Protease with a molar ratio of 1:100 (C_fusion protein_ : C_PreScission Protease_). The digest was performed for 16h at 8°C with gentle agitation.

To separate the ENTH-domain from the cleaved GST-tag an anion exchange chromatography was performed by using HiTrap™ Q HP, 5 ml (GE Healthcare Life Science, Chalfont St Giles, UK). The protein suspension was loaded onto the column by using anion exchange low salt buffer (50 mM NaCl, 50 mM HEPES/NaOH, pH 8.0) with a flow of 1 ml/min at 8°C. The free ENTH-domain eluted during the washing with the same buffer (3 ml/min, 8°C) while GST remained until using anion exchange high salt buffer (1 M NaCl, 50 mM HEPES/NaOH, pH 8.0). A size-exclusion chromatography HiLoad™ 16/600 Superdex™ 75 pg (GE Healthcare Life Science, Chalfont St Giles, UK) was used in the final step to ensure the purity of the ENTH-domain. The ENTH-domain was eluted with HEPES buffer (or PBS buffer if it was labeled afterwards) at 1 ml/min and 8°C. The proteins were directly used afterwards or stored at −80°C.

#### Visualization of LUV deformation by electron microscopy

Large unilamellar vesicles were prepared as described before ^31,32^. Briefly, L-α-phosphatidylcholin (PC), L-α-phosphatidylethanolamine (PE), L-α-phosphatidylserine (PS) and L-α-Phosphatidylinositol-4,5-bisphosphate (PIP_2_) were obtained from Avanti Polar Lipids (Alabaster, AL).

To observe the protein induced deformation of LUVs (as described before ^33,34^), liposomes (0.2 mg/ml) were incubated with ENTH domain or it’s mutants (15 μM) at 30°C for 3 h. The samples were subsequently diluted to 0.2 mg/ml of liposomes with HEPES buffer (200 mM NaCl, 10 mM HEPES/NaOH, pH 7.4) and 5 μl of the suspension was then transferred onto a formvar carbon coated copper grid (Agar Scientific Ltd., Essex, UK) and incubated for 1 min at room temperature. The suspension was removed and the grid was set onto a droplet (50 μl) of 3% uranyl acetate for negative staining.

Electron microscopic visualization was performed with a JEOL JEM 1011 transmission electron microscope (JEOL Ltd., Akishima, Japan) and a Gatan Orius 1000 CCS detector (Gatan Inc., Pleasanton, USA).

#### Binding and membrane deformation dynamics on GUVs

GUVs were prepared as previously described with 0.5 % of the fluorescently labeled lipid Atto647N PtdEnt (ATTO-TEC GmbH, Siegen, Germany). 50 μl of the GUV suspension was carefully transferred into the microscopy chamber Nunc^®^ Lab-Tek^®^ II chambered coverglass (Thermo Fisher Scientific Inc., Waltham, USA), that was coated with lipid-free bovine serum albumin (BSA, Sigma-Aldrich Crop., St. Louis, USA) and filled with 250 μl of GUV buffer (192 mM NaCl, 10 mM HEPES/NaOH, pH 7.4).

The fluorescence spinning disc confocal microscopy was performed at 25°C with a Nikon Eclipse Ti microscope (Nikon Instruments K.K., Tokyo, Japan), a PerkinElmer UltraVIEW VoX system (Perkin Elmer, Waltham, USA) with a spinning disk confocal scan head CSU-X1 (Yokogawa Denki K. K., Tokyo, Japan) and an Electron Multiplying CCD Camera C9100 (Hamamatsu Photonics K.K., Hamamatsu, Japan). The data was acquired with the software Volocity^®^ 6.3 (PerkinElmer Corp., Waltham, USA). Proteins that were labeled with Atto488 maleimide (ATTO-TEC GmbH, Siegen, Germany) were titrated into the GUV suspension by 10 μM steps. The fluorophores were excited at the optimal wavelength and detected at the optimal emission wavelength according to the manufacturer (ATTO-TEC GmbH, Siegen, Germany). All fluorophores were excited at low laser powers (5% to 8%) and at fast exposure times (50 – 120 msec). Images were taken with a speed of 30 frames per minute for 30 min.

#### Secondary structure analysis by CD spectroscopy

Circular dichroism (CD) spectroscopy was used to compare the folding of the wildtype ENTH domain and the ENTH domain mutant R114A by using a Chirascan CD Spectrometer (Applied Photophysics Ltd., London, UK). Proteins were transferred into a 10 mM sodium potassium phosphate buffer pH 7.4 with a concentration of 0.1 mg/ml. The measurements were carried out at 25°C between 180 nm and 260 nm with 1 nm step sizes and an accumulation time of 5 sec. The mean residue weight ellipticity was calculated according to Kelly S. M. et al. ^188^ with the following formula:

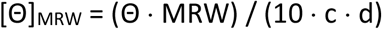

The molar ellipticity *Θ* and the mean residue weight (MRW) are dependent on the optical path length *d* and the protein concentration *c*.

#### Co-sedimentation assay (Spin assay)

The co-sedimentation assay (spin assay) was used for two purposes:

1. To detect protein binding to LUVs by co-sedimentaion
2. To detect if small vesicles were generated due to membrane deformation. The sedimentation of spherical particles in aqueous solution is described by Stokes’s law.

The sedimentation behavior of a liposome is more dependent on the size than on the density and it is possible to separate SUVs from LUVs by centrifugation.

LUVs were prepared as described before, diluted to 2 mg/ml and mixed with 20 μM purified proteins. After incubating the sample for 3 h at 30°C, 150 μl of the sample was ultracentrifuged at 22°C and 205000 × *g* for 19 min (TLA 100.3, Beckman Coulter Inc., Fullerton, California). The experiments were performed in HEPES buffer. After separating the supernatant from the pellet both were used for SDS-PAGE analysis.

#### LUV uniformity analysis by DLS

The LUV suspension (with and without protein) was diluted to 0.2 mg/ml and set into the measurement chamber of a Zetasizer Nano-S (Malvern Instruments Ltd., Worcestershire, UK). The data was acquired by three independent measurements, each one consisting of 21 repetitions at 25°C and using the Zetasizer Software 7.01 (Malvern Instruments Ltd., Worcestershire, UK).

#### FRET assay

FRET measurements were performed to detect the oligomerization of ENTH domains and the interaction between these proteins and specific lipids. Atto488 labeled ENTH domains were used as donor and Atto532 coupled to DOPS, DOPE (ATTO-TEC GmbH, Siegen, Germany), or ENTH domains were used as acceptor. The donor was excited at 493 nm and the acceptor emission maximum is at 552 nm. A total concentration of 5 and 15 μM labeled ENTH domain added into a 2 mg/ml lipid suspension. The FRET experiments were performed in a F-7000 Fluorescence Spectrophotometer (Hitachi K. K., Tokio, Japan) and the emission spectra were recorded between 500 – 700 nm.

#### Fluorescence labeling of ENTH domains

The ENTH WT (C96A A155C) domain and the ENTH R114A (C96A A155C) were labeled with Atto488 maleimide (or Atto532 maleimide) (ATTO-TEC GmbH, Siegen, Germany) by cysteine modification. The substitution reaction took place in PBS (136 mM NaCl, 2.7 mM KCl, 8 mM Na_2_HPO_4_, 1.8 mM K_2_HPO_4_, pH 7.4) with a molar access of the fluorophore of 1.4 in respect to the protein. The substitution was performed at 4°C with gentle agitation on a tumbling shaker during 16 h.

Proteins were finally separated from free fluorophores by size exclusion chromatography with a PD MiniTrap G-25 prepacked columns (GE Healthcare Life Science, Chalfont St Giles, UK). Finally, the degree of labeling (DoL) of the proteins was determined spectrophotometrically by measuring the UV-Vis (NanoDrop 2000, Thermo Fisher Scientific Inc., Waltham, USA). For this purpose, the following equation was used:

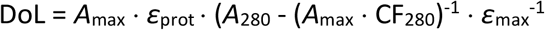

A_max_ is the absorption at the characteristic wavelength of the fluorophore, ε_max_ is the molar extinction coefficient of the fluorophore, *A*_280_ is the absorption of the protein at 280 nm, *ε*_prot_ is the molar extinction coefficient of the labeled protein and CF_280_ is the correction factor of the fluorophore.

### RIfS and AFM

#### Vesicle preparation

1-Palmitoyl-2-oleoyl-*sn*-glycero-3-phosphocholine (POPC), 1-Palmitoyl-2-oleoyl-*sn*-glycero-3-phospho-L-serine (POPS), 1,2-dioleoyl-*sn*-glycero-3-phosphocholine (DOPC), 1,2-dioleoyl-*sn*-glycero-3-phosphoethanolamine (DOPE), 1,2-dioleoyl-*sn*-glycero-3-phospho-L-serine (DOPS) and L-α-Phosphatidylinositol-4,5-bisphosphate (PIP_2_) were obtained from Avanti Polar Lipids (Alabaster, AL). Sulforhodamine-1,2-dihexanoyl-sn-glycero-3-phosphoethanolamine (Texas Red DHPE) was purchased from Sigma-Aldrich (Taufkirchen, Germany) and silicon wafers from Silicon Materials (Kaufering, Germany).

Stock solution of lipids were prepared in chloroform at a concentration of 2-10 mg/mL. For PI(4,5)P_2_ a mixture of chloroform/methanol was used. The lipid stock solutions and fluorophores were added to a test tube filled with 400 μL chloroform and 30 μL methanol, obtaining the desired molar ratio. After removing the solvent in a nitrogen flush the lipid film was further dried under vacuum at 30° C for 3 h and stored at 4° C until use.

To obtain small unilamellar vesicles (SUVs), the lipid films (0.4 mg) were rehydrated with 0.5 mL citrate buffer (20 mM Na-citrate, 50 mM KCl, 0.1 mM EDTA, 0.1 mM NaN_3_, pH 4.8) for 30 min, followed by vortexing (3 × 30 s in a 5 min interval). By sonication of the lipid suspension for 30 min (60% power, Sonopuls HD2070, Bandelin; Berlin, Germany) the SUV suspension was gained.

#### Preparation of Solid Supported Lipid Membranes

For the AFM and RIfS experiments solid supported lipid membranes (SLBs) were prepared on silicon dioxide. Silicon wafers with 100 nm SiO_2_ (AFM) and 5000 nm SiO_2_ (RIfS) were cut into 0.8 × 1.9 cm^2^ rectangles and treated in an aqueous solution of NH_3_ and H_2_O_2_ (H_2_O/NH_3_/H_2_O_2_ 5:1:1) for 30 min at 70 °C to obtain a hydrophilic surface. Further hydrophilization was reached by oxygen plasma treatment (30 s, 60% power).

The AFM substrates were fixed in a Teflon chamber and incubated with SUVs consisting of DOPC/DOPE/PI(4,5)P_2_/Texas Red DHPE 44.9:30:5:0.1 and DOPC/DOPE/DOPS/PI(4,5)P_2_/Texas Red DHPE 64.9:30:20:5:0.1 for 60 min. Rinsing with citrate-buffer and PBS (1.5 mM KH_2_PO_4_, 8.1 mM Na_2_HPO_4_, 2.7 mM KCl, 136.9 mM NaCl, pH 7.4) to removed remaining lipid material. The time-resolved SLB formation via RIfS is described below.

#### Reflectometric Interference Spectroscopy (RIfS)

The formation of SLBs and the protein adsorption were monitored using the fiber optic spectrometer SD2000 (Ocean Optics, Dunedin, USA) and the miniature spectrometer flame (Ocean Optics, Dunedin, USA). The experimental setup was already described previously ^35^. The recorded spectra (every 2 s) were evaluated with a MATLAB routine. The hydrophilized substrates were implemented in a chamber and rinsed with ultrapure water and citrate buffer. SUVs of desired lipid compositions were added in a circle. After reaching a constant optical thickness, indicating the successful spreading of a bilayer, the substrates were rinsed with PBS to remove remaining lipid material. Passivation with BSA solution (1 mg/mL in PBS) prevented non-specific interactions. Afterwards the system again was rinsed with PBS. Then 0.1-5 μM ENTH WT and ENTH R114A were added until a plateau was reached. Rinsing with PBS resulted in ENTH desorption.

#### Atomic force microscopy (AFM)

To investigate the protein coverage and protein height, atomic force micrographs of the SLB surface before and after protein addition were taken. ENTH^WT^ and ENTH^R114A^ (1 μM) were incubated for 2 h at RT. The solution was mixed with a stirring bar to ensure homogenous distribution of the protein. Afterwards 10 × 10 μm^2^ and 1 × 1 μm^2^ areas of the substrates were imaged in contact mode with BL-AC40TS-C2 cantilevers (*f* = 85.4-139.1 kHz, *k* = 0.03-0.12 N m^−1^, Olympus, Tokio, Japan). The measurements were performed using a JPK Nanowizard 4 (JPK Instruments, Berlin, Germany) equipped with a CCD camera (Fire Wire CCD color camera, The Imaging Source, NC, USA). The protein height and density were calculated with a MATLAB routine.

### Cell Biology

#### Plasmids

pEGFP-N1 (Clonetech), pEGFPC1-epsin 1 ^36^. pEGFPC1-epsin 1^R114A^ was prepared by QuikChange II Site-directed mutagenesis kit (Agilent) using the pEGFPC1-epsin 1 as a template. Suppression (knock-down) of epsin 1, 2 and 3 expression in HeLa (ATCC^®^ CCL-2™) cells by RNA interference approach was performed as described in ^37^. This approach was taken due to the redundant functions of epsin 1, 2 and 3 (Boucrot et al., 2012). Here, HeLa cells were electroporated with scrambled or three different DsiRNAs (Integrated DNA Technologies, Inc. (IDT), Coralville, Iowa, Epsin1: hs.Ri.EPN1.13.3; Epsin2: hs.Ri.EPN2.13.1; Epsin3: hs.Ri.EPN3.13.3) two times (with a 24 h interval) and plated on the poly-L-lysine coated glass coverslips (Marienfeld, Cat No. 0117580; Ø18 mm) in the DMEM media (Gibco) containing 10% fetal bovine serum (FBS) and penicillin/streptomycin (Gibco). The efficiency of KD was inspected by quantitative PCR (qPCR; Figure S4A), as in Boucrot et al. (2012), since the specific antibodies for three epsin proteins (1, 2 and 3) are not commercially available. After 20 h on coated coverslips, cells were transfected with either EGFP, epsin 1^WT^-EGFP or epsin 1^R114A^-EGFP using Fugene 6 transfection reagent (Promega). For the fluorescent uptake assays, 28-32h after transfection cells were serum-starved for 1 h at 37°C in pre-warmed DMEM containing 20 mM HEPES, pH 7.4, and 1% BSA, and then incubated with biotinylated Epidermal Growth Factor (EGF) complexed to Texas Red™ Streptavidin (ThermoFischer/Molecular Probes Cat. No. E3480; referred to as EGF-Texas Red in the text and figures; 2 μg/mL) or Alexa Fluor 548–transferrin (ThermoFischer/Molecular probes Cat. No. T23364; referred to as transferrin-A548 in the text and figures; 50 μg/mL) on ice for 30 min. Subsequently, cells were washed once in ice-cold PBS and then placed in pre-warmed DMEM media for indicated time at 37°C. After incubation, cells were placed on ice and incubated in cold acidic PBS (pH 4.5) for 5 min on ice to remove fluorescent EGF or transferrin from the cell surface. The cells were then washed two times with ice-cold PBS (pH 7.4), fixed and processed for immunofluorescence. In experiments with clathrin-mediated inhibitor Pitstop-2 (Abcam, Cat. No, ab120687), the inhibitor was added 30 min before the experiment was started, and kept throughout the transferrin-A548 uptake. Images were captured with a commercial confocal Zeiss 800 Airyscan microscope setup (Carl Zeiss Inc.). The acquired images were analyzed using Metamorph software 6.1. (Molecular Devices) and displayed using Sigma Plot 12.5 (Systat Software, Inc.) and Adobe Photoshop 10 (Adobe). The background is subtracted for each cell. Alternatively, epsin 1,2,3-KD HeLa cells were plated after electroporation on poly-L-lysine-coated plastic petri dishes (Sarstadt Ø18 mm) in the DMEM media (Gibco) containing 10% fetal bovine serum (FBS) for 20-22h, and transfected with either EGFP, epsin 1 WT-EGFP or epsin 1^R114A^-EGFP as before. 28-34h after transfection, cells were serum-starved for 1 h at 37°C in pre-warmed DMEM containing 20 mM HEPES, pH 7.4, and 1% bovine serum albumin (BSA) and then incubated with biotinylated Epidermal Growth Factor (EGF) complexed to Texas Red™ Streptavidin (ThermoFischer/Molecular Probes Cat. No. E3480; referred to as EGF-Texas Red in the text and figures; 2 μg/mL) or Alexa Fluor 548–transferrin (ThermoFischer/Molecular probes Cat. No. T23364; referred to as transferrin-A548 in the text and figures; 50 μg/mL) on ice for 30 min, washed once in ice-cold PBS and then placed in pre-warmed DMEM media for indicated time at 37°C. After incubation, cells were placed on ice and incubated in cold acidic PBS (pH 4.5) for 5 min on ice to remove fluorescent EGF or transferrin from the cell surface. Cells were then harvested from the plates and processed for flow cytometry. Fluorescence-activated cell sorting (FACS) was done by Sony Cell Sorter SH800S. FACS data were analyzed by FlowJo software and plotted in Prism (GraphPad).

### NMR spectroscopy

NMR samples containing ENTH and its variant R119 were packed in a 3.2 mm rotor with DSS added externally for temperature calibration. Experiments were recorded on a 850MHz wide-bore spectrometer equipped with 3.2 mm triple resonance probe (Bruker Biospin). All ssNMR experiments were recorded at 5 °C. 90° pulse widths of 3μs for ^1^H, 5μs for ^13^C, 7μs for ^15^N and decoupling strength of ~ 70-80 kHz was used for CP and C90 experiment. Spectra were processed and analysed in Topspin. The samples were spun at 8 kHz for INEPT experiment, 11kHz for CP and C90 experiment.

